# Single-cell analysis of human lung epithelia reveals concomitant expression of the SARS-CoV-2 receptor ACE2 with multiple virus receptors and scavengers in alveolar type II cells

**DOI:** 10.1101/2020.04.16.045617

**Authors:** Guangchun Han, Ansam Sinjab, Warapen Treekitkarnmongkol, Patrick Brennan, Kieko Hara, Kyle Chang, Elena Bogatenkova, Beatriz Sanchez-Espiridion, Carmen Behrens, Boning Gao, Luc Girard, Jianjun Zhang, Boris Sepesi, Tina Cascone, Lauren Byers, Don L. Gibbons, Jichao Chen, Seyed Javad Moghaddam, Edwin J. Ostrin, Junya Fujimoto, Jerry Shay, John V. Heymach, John D. Minna, Steven Dubinett, Paul A. Scheet, Ignacio I. Wistuba, Edward Hill, Shannon Telesco, Christopher Stevenson, Avrum E. Spira, Linghua Wang, Humam Kadara

## Abstract

The novel coronavirus SARS-CoV-2 was identified as the causative agent of the ongoing pandemic COVID 19. COVID-19-associated deaths are mainly attributed to severe pneumonia and respiratory failure. Recent work demonstrated that SARS-CoV-2 binds to angiotensin converting enzyme 2 (ACE2) in the lung. To better understand *ACE2* abundance and expression patterns in the lung we interrogated our in-house single-cell RNA-sequencing dataset containing 70,085 EPCAM+ lung epithelial cells from paired normal and lung adenocarcinoma tissues. Transcriptomic analysis revealed a diverse repertoire of airway lineages that included alveolar type I and II, bronchioalveolar, club/secretory, quiescent and proliferating basal, ciliated and malignant cells as well as rare populations such as ionocytes. While the fraction of lung epithelial cells expressing *ACE2* was low (1.7% overall), alveolar type II (AT2, 2.2% *ACE2*+) cells exhibited highest levels of *ACE2* expression among all cell subsets. Further analysis of the AT2 compartment (n = 27,235 cells) revealed a number of genes co-expressed with *ACE2* that are important for lung pathobiology including those associated with chronic obstructive pulmonary disease (COPD; *HHIP*), pneumonia and infection (*FGG* and *C4BPA*) as well as malarial/bacterial (*CD36*) and viral (*DMBT1*) scavenging which, for the most part, were increased in smoker versus light or non-smoker cells. Notably, *DMBT1* was highly expressed in AT2 cells relative to other lung epithelial subsets and its expression positively correlated with *ACE2*. We describe a population of *ACE2*-positive AT2 cells that co-express pathogen (including viral) receptors (e.g. *DMBT1*) with crucial roles in host defense thus comprising plausible phenotypic targets for treatment of COVID-19.

## INTRODUCTION

In late December 2019, an outbreak of lung pneumonia initially of unknown cause was reported in China ^1^. This emerging disease, termed coronavirus disease 2019 (COVID-19), was soon thereafter attributed to infection with the novel zoonotically-transmitted coronavirus severe acute respiratory syndrome coronavirus 2 (SARS-CoV-2) ^2^. On March 11, 2020, COVID-19 was declared a rapidly spreading global pandemic by the world health organization. As of April 12, SARS-CoV-2 has infected more than 1.8 million people worldwide resulting in over 110,000 deaths ^3^.

Upon SARS-CoV-2 infection, the clinical presentation of COVID-19 is diverse, ranging from asymptomatic infection and mild upper respiratory illness to pneumonia, acute respiratory distress syndrome (ARDS), respiratory failure and death ^1^. Recent clinical reports have suggested that old age and comorbidities such as cardiovascular disease, chronic inflammatory lung disease and cancer are risk factors for COVID-19-associated severe pneumonia and death ^1^. Recent analyses reported that COVID-19 patients who were smokers required more aggressive clinical interventions and that smoking possibly elevates the risk of COVID-19 associated pneumonia and death ^4, 5^. These findings suggest that limiting inflammatory responses by intercepting pathways involved pulmonary (and perhaps cardiac) pathobiology may represent viable strategies to improve COVID-19 patient outcomes.

Recent molecular studies have demonstrated that SARS-CoV-2 infects airway cells by binding of its spike protein to angiotensin converting enzyme 2 (ACE2), the same receptor used by SARS-CoV ^6, 7, 8, 9^. While SARS-CoV and SARS-CoV-2 spike proteins exhibit high degrees of homology, recent studies have demonstrated that the novel coronavirus binds ACE2 more efficiently than the original SARS-CoV virus ^10, 11, 12, 13^ Expression of ACE2 in the lung was suggested to implicate lung epithelial cells as routes of entry and, thus, targets for SARS-CoV-2 infection. ACE2 is a carboxypeptidase that limits activation of the renin-angiotensin system (RAS), a hormonal system that plays major roles in cardiovascular and pulmonary physiology and pathology ^14^. ACE2 has been shown to protect from lung injury mediated by SARS-CoV infection which was also alleviated by targeting the RAS ^15^. A large body of literature suggests that the ACE2-mediated RAS axis plays important roles in lung function ^14^, susceptibility to the SARS virus ^8^, and inhibition of angiogenesis in lung cancer ^16^. However, the role of key pathways in lung pathobiology and inflammation in the pathogenesis of COVID-19, particularly in high risk individuals, remains unexplored.

Motivated by the current and unprecedented global health crisis, we leveraged our ongoing lung single-cell RNA-sequencing (scRNA-seq) efforts to interrogate the expression of the SARS-CoV-2 receptor *ACE2* in a unique cohort of 70,085 lung epithelial cells from tumor and matched uninvolved normal lung tissues from five lung adenocarcinoma (LUAD) patients. Among all lung epithelial subsets, we found that *ACE2* expression was higher in uninvolved AT2 cells and malignant epithelial cells. We interrogated single-cell gene co-expression profiles and found that *ACE2*-epressing AT2 cells concurrently expressed other genes with important yet understudied roles in lung pathobiology including fibrosis, modulation of chronic inflammation and innate host-defense mechanisms by additional viral receptors and scavengers, particularly among cells from smokers. Our single-cell analyses highlight additional possible receptors that may play a role in SARS-CoV-2 infection in the lung epithelium and, thus, are plausible targets that can be repurposed for clinical management of COVID-19.

## MATERIALS AND METHODS

### Derivation of single cells from lung tissues

Two to three uninvolved normal lung tissues per patient and LUADs (n = 19 samples) were collected from surgically resected specimens from five patients with early-stage disease (stages I-IIIA) (**Table 1**). All patients were evaluated at the University of Texas MD Anderson Cancer Center (MD Anderson) and had provided informed consents under approved institutional review board protocols. Fresh tissues were collected in RPMI medium supplemented with 2% fetal bovine serum (FBS) and maintained on ice for immediate processing. Tissues were placed in a cell culture dish containing Hank’s balanced salt solution (HBSS) on ice and extra-pulmonary airways and connective tissue was removed with scissors. Samples were transferred to a new dish on ice and minced into tiny pieces (approximately 1mm^3^) followed by enzymatic digestion using a cocktail containing Collagenase A (10103578001, Sigma Aldrich), Collagenase IV (NC9836075, Thermo Fisher Scientific), DNase I, (11284932001, Sigma Aldrich), Dispase II (4942078001, Sigma Aldrich), Elastase 1 (NC9301601, Thermo Fisher Scientific) and Pronase (10165921001, Sigma Aldrich). Samples were incubated in a 37 °C oven for 10 minutes with gentle rotation and then incubated for an additional 10 minutes. Samples were then filtered through 70 μm strainers (Miltenyi biotech, 130-098-462) and washed with ice-cold HBSS. Filtrates were then centrifuged and re-suspended in ice-cold ACK lysis buffer (A1049201, Thermo Fisher Scientific) for red blood cell (RBC) lysis. Following RBC lysis, samples were centrifuged and re-suspended in ice-cold FBS, filtered (using 40 μm FlowMi tip filters; H13680-0040, Millipore). Cells were enumerated and examined for viability by Trypan blue exclusion analysis. Cells were immediately frozen in FBS containing 10% dimethyl sulfoxide and cryopreserved.

**Table 1.** Clinicopathological variables of the LUAD patients analyzed by scRNA-seq.

| **Patient** | **Stage** | **Age** | **Smoking status** | **Sex** | **Ethnicity** |
| --- | --- | --- | --- | --- | --- |
| P1 | IIIA | 70 | Non-smoker | Female | White or Caucasian |
| P2 | IB | 73 | Former | Female | White or Caucasian |
| P3 | IIIA | 58 | Light former* | Female | White or Caucasian |
| P4 | IIB | 64 | Former | Female | White or Caucasian |
| P5 | IB | 71 | Non-smoker | Male | White or Caucasian |

\*light smoker (less than five pack-years of smoking)

### Sorting and enrichment of viable lung EPCAM+ epithelial singlets

At the time of processing, cryopreserved single cells from patient 1 were rapidly thawed, washed, and stained with SYTOX Blue viability dye (S34857, Life technologies), and processed on a FACS Aria II instrument. Cells from patients 2 through 5 were stained following washing with anti-EPCAM-PE (347198, BD Biosciences;1:50 dilution in ice-cold phosphate-buffered saline, PBS, containing 2% FBS) for 30 minutes. EPCAM-stained cells were then washed, filtered using 40 μm tip filters, stained with SYTOX Blue and processed on a FACS Aria II instrument. Doublets and dead cells were eliminated, and viable (SYTOX-negative) EPCAM-positive (EPCAM+, epithelial) singlets were collected in PBS containing 2% FBS. Cells were washed again to eliminate ambient RNA, and a sample was taken for counting by Trypan Blue exclusion (T8154, Sigma Aldrich) before loading on 10X Chromium microfluidic chips.

### Preparation of single-cell 5’ gene expression libraries

Up to 10,000 cells per sample were partitioned into nanoliter-scale Gel beads-in-emulsion (GEMs) using Chromium Next GEM Single Cell 5’ Gel Bead Kit v1.1 (1000169, 10X Genomics) and by loading onto Chromium Next GEM Chips G (1000127, 10X Genomics). GEMs were then recovered to construct single-cell 5’ gene expression libraries using the Chromium Next GEM Single Cell 5’ Library kit (1000166, 10X Genomics) according to the manufacturer’s protocol. Briefly, recovered barcoded GEMs were broken and pooled, followed by magnetic bead clean-up (Dynabeads MyOne Silane, 37002D, Thermo Fisher Scientific). 10X-barcoded full-length cDNA was then amplified via PCR and analyzed using Bioanalyzer High Sensitivity DNA kit (5067-4626, Agilent). Up to 50 ng of cDNA was carried over to construct gene expression libraries and was enzymatically fragmented and size-selected to optimize the cDNA amplicon size prior to 5’ gene expression library construction. Further, samples were subject to end-repair, A-tailing, adaptor ligation, and sample index PCR using Single Index Kit T Set A (2000240, 10X Genomics) to generate Illumina-ready barcoded gene expression libraries. Library quality and yield was measured using Bioanalyzer High Sensitivity DNA kit (5067-4626, Agilent) and Qubit dsDNA High Sensitivity Assay Kit (Q32854, Thermo Fisher Scientific). Indexed libraries were normalized before pooling by adjusting for the ratio of the targeted cells per library as well as individual library concentration and pooled to a final concentration of 10 nM. Library pools were then denatured and diluted as recommended for sequencing on the Illumina NovaSeq 6000 platform.

### scRNA-seq data analysis

Single-cell decoding was performed using available computational framework. Raw scRNA-seq data was pre-processed, demultiplexed, aligned to human GRCh38 reference and feature-barcodes generated using CellRanger (10X Genomics, version 3.0.2). Detailed quality control metrics were generated and evaluated. Genes detected in < 3 cells and cells where < 200 genes had non-zero counts were filtered out and excluded from subsequent analysis. Low quality cells where >15% of the read counts originated from the mitochondrial genome were also discarded. In addition, cells with >6,000 detected genes were discarded to remove likely doublet or multiplet captures. Possible batch effects were evaluated and corrected using Harmony ^17^. Raw unique molecular identifier (UMI) counts were log normalized and used for principal component analysis (PCA). We applied Seurat ^18^ for clustering and UMAP ^19^ for dimensional reduction to perform unsupervised analysis of the high-dimensional data into distinct cell clusters. EPCAM+ cells were partitioned into major airway lineage clusters, followed by subclustering within each compartment/lineage to identify subpopulations. Differentially expressed genes (DEGs) for each cell cluster were identified using the *FindAllMarkers* function in Seurat R package. The top 50 most significant DEGs were examined and an integrative approach was used to determine cell types and states.

In addition to the bioinformatics approaches described above, all other statistical analysis was performed using R v3.6.0. Analysis of differences in genes between groups (e.g., between *ACE2*-postive vs. *ACE2*-negative; smoker vs. non/light smoker) were calculated using the *FindMarkers* function in R. Pseudo-bulk gene expression values for defined cell clusters were calculated by taking mean expression of each gene across all cells in a specific cluster. Deconvolution of AT2 cell abundance was performed on the TCGA lung adenocarcinoma cohort by incorporating AT2 signature genes defined by our study into MCP-counter, an R package that permits the quantification of the absolute abundance of immune and stromal cell populations in heterogeneous tissues from transcriptome data ^20^. All statistical significance testing was two-sided, and results were considered statistically significant at p-value < 0.05. The Benjamini-Hochberg method was applied to control the false discovery rate (FDR) in multiple comparisons (e.g. DEG analysis), and calculate adjusted p-values (q-values).

## RESULTS

### Single-cell decoding of normal and malignant lung epithelial cells

We performed single-cell analysis of normal lung tissues and matched treatment-naïve early-stage lung adenocarcinomas (LUADs) from five patients using droplet based scRNA-seq. We first processed two normal lung tissues and a LUAD from patient 1 which resulted in 15,370 cells that were retained for analysis. In line with studies of other organs ^21^, we noted that the fraction of epithelial (*EPCAM*+) cells attained by unbiased analysis of lung tissues was limited (∼4%, n = 624 cells) (**Table 2**). To better capture lung epithelial cells, including their heterogeneity, we performed single-cell analysis of fluorescent-activated cell sorting (FACS) enriched EPCAM+ cells from three normal lung tissues (per patient) and LUADs from four additional patients. Using this strategy, we were able to capture on average 20-fold more lung epithelial cells (**Table 2**). Following quality control and filtering, we retained 70,085 lung epithelial cells from the 19 samples (5 tumor samples, n = 13,134 cells; 14 normal lung tissues, n = 56,951 cells) (**Fig. 1a**). We achieved on average 146,574 reads, 9,202 unique molecular identifiers and 2,407 genes per cell (**Table 2**). Unsupervised clustering analysis identified a diverse repertoire of airway lineages which were divided into 10 cell clusters with distinct transcriptomic features (e.g., expression of lineage-specific markers) (**Fig. 1a**). These cell clusters covered major lineages of lung epithelium such as alveolar type I (AT1; n = 10,775; *AGER, EMP2, CAV1*), AT2 (n = 27,235; *SFTPC, SFTPB, PGC*), club and secretory (n = 4,625; *SCGB1A1, SCGB3A1, SCGB3A2*), ciliated (n = 3,247; *FOXJ1, CAPS, PIFO*) and basal cells (n = 5,119; *KRT17, KRT15, TP63*) including those with proliferative properties (n = 449; *MKI67, TOP2A*) (**Fig. 1c** and **Fig. S1**). One cluster included bronchioalveolar cells that were enriched for markers of both AT2 and club lineages (**Fig. 1b**). We also identified alveolar transitory or progenitor cells which expressed both AT2 and AT1 markers including *HOPX* as well as *KRT7* (**Fig. 1c**) in line with previous reports ^22, 23^. Clustering analysis also revealed a rare cell population that exclusively expressed *ASCL3* and *FOXI* (**Fig. 1c** and **Fig. S1**) characteristic of pulmonary ionocytes which were recently described by single-cell analysis ^24, 25^. Of note, most cells from the LUAD specimens clustered distinctly from cells of uninvolved lung tissues and exhibited the highest expression of the tumor marker *CEACAM5* and mixed lineage genes (**Fig. 1b-c**) thus representing diverse lung malignant cells in line with recent studies ^26^.

**Table 2.**
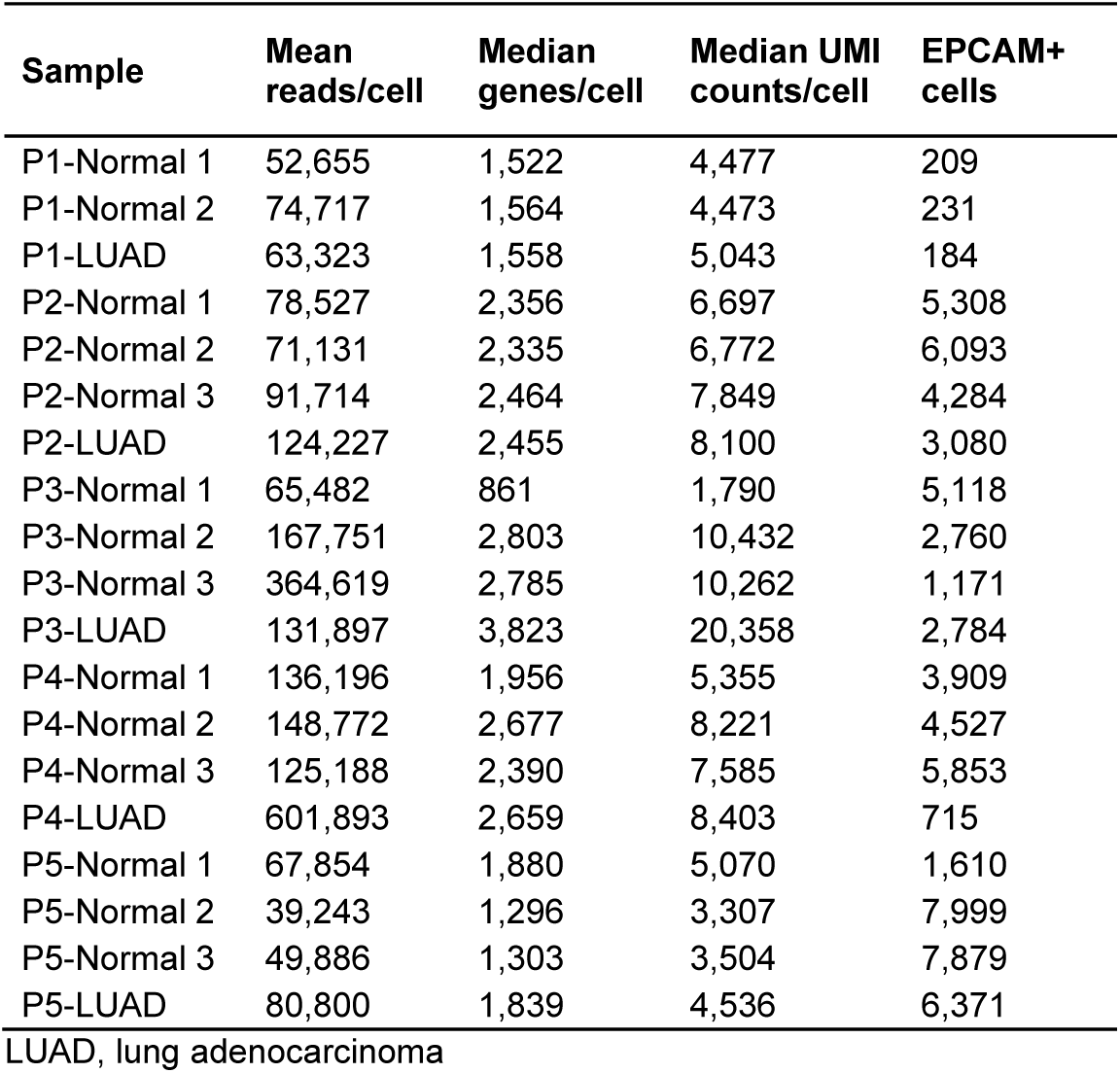
Number of EPCAM+ cells analyzed in normal lung and LUAD tissues.

| **Sample** | **Mean reads/cell** | **Median genes/cell** | **Median UMI counts/cell** | **EPCAM+ cells** |
| --- | --- | --- | --- | --- |
| P1-Normal 1 | 52,655 | 1,522 | 4,477 | 209 |
| P1-Normal 2 | 74,717 | 1,564 | 4,473 | 231 |
| P1-LUAD | 63,323 | 1,558 | 5,043 | 184 |
| P2-Normal 1 | 78,527 | 2,356 | 6,697 | 5,308 |
| P2-Normal 2 | 71,131 | 2,335 | 6,772 | 6,093 |
| P2-Normal 3 | 91,714 | 2,464 | 7,849 | 4,284 |
| P2-LUAD | 124,227 | 2,455 | 8,100 | 3,080 |
| P3-Normal 1 | 65,482 | 861 | 1,790 | 5,118 |
| P3-Normal 2 | 167,751 | 2,803 | 10,432 | 2,760 |
| P3-Normal 3 | 364,619 | 2,785 | 10,262 | 1,171 |
| P3-LUAD | 131,897 | 3,823 | 20,358 | 2,784 |
| P4-Normal 1 | 136,196 | 1,956 | 5,355 | 3,909 |
| P4-Normal 2 | 148,772 | 2,677 | 8,221 | 4,527 |
| P4-Normal 3 | 125,188 | 2,390 | 7,585 | 5,853 |
| P4-LUAD | 601,893 | 2,659 | 8,403 | 715 |
| P5-Normal 1 | 67,854 | 1,880 | 5,070 | 1,610 |
| P5-Normal 2 | 39,243 | 1,296 | 3,307 | 7,999 |
| P5-Normal 3 | 49,886 | 1,303 | 3,504 | 7,879 |
| P5-LUAD | 80,800 | 1,839 | 4,536 | 6,371 |

LUAD, lung adenocarcinoma

**Figure 1:**
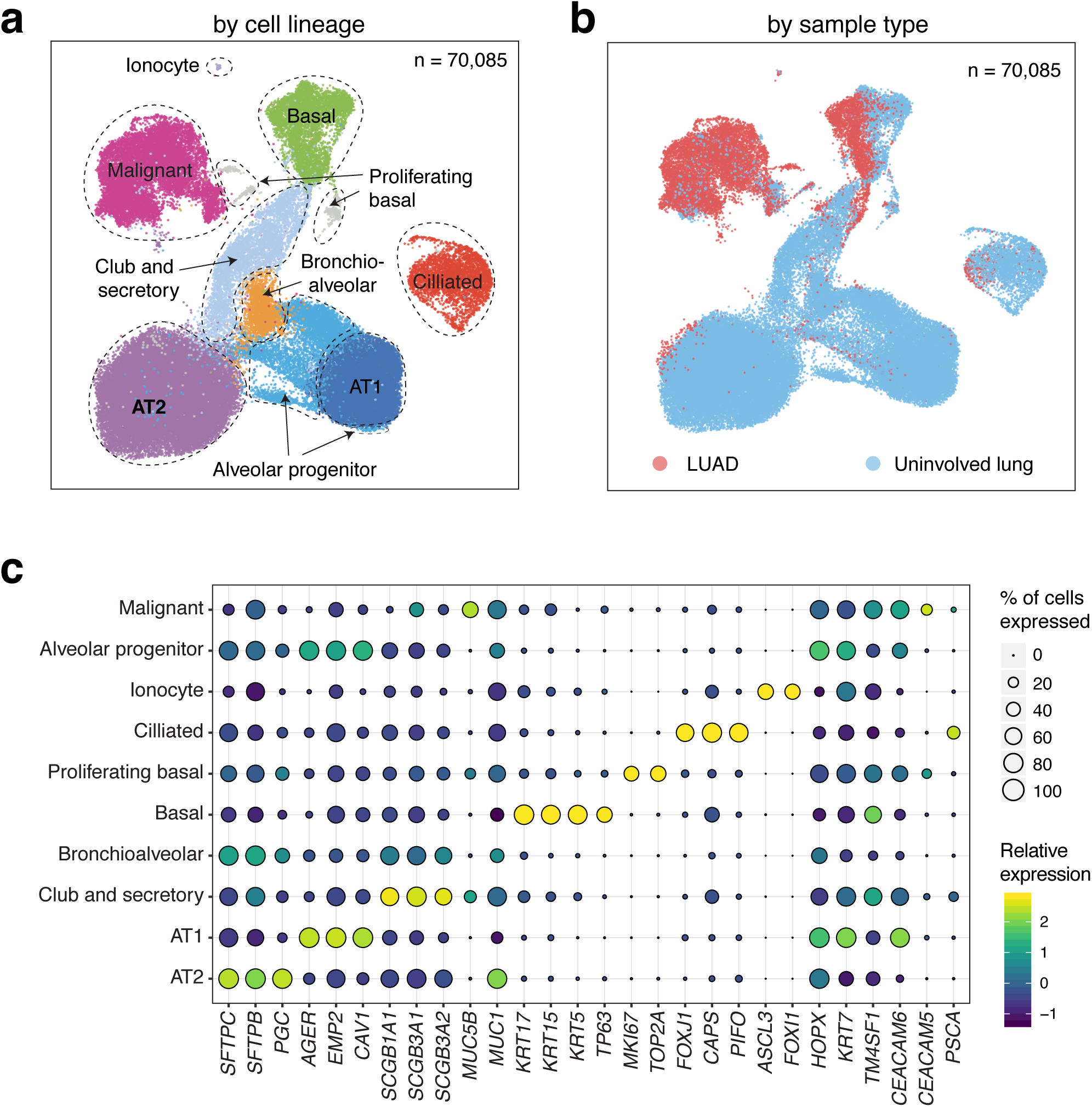
Overview of 70,085 EPCAM+ lung epithelial cells analyzed by scRNA-seq. UMAP plots with cells labeled by lineage (**a**) and sample type (**b**). AT1: alveolar type I cells; AT2: alveolar type II cells; LUAD: lung adenocarcinoma. **c**) Bubble plots showing cell fractions with scaled expression of airway lineage markers for the identified lung epithelial cell clusters.

### Abundance and expression patterns of the SARS-CoV-2 receptor *ACE2* in lung epithelium

The ongoing COVID-19 pandemic caused by infection with the novel coronavirus SARS-CoV-2 prompted us to leverage our lung scRNA-seq dataset and interrogate expression patterns of the SARS-Cov-2 receptor *ACE2* in lung epithelial cells. We found that the fraction of *ACE2*-expressing cells among all lung epithelial cells was low (n= 1,208, 1.7%) (**Fig. 2a**). The highest fractions of *ACE2*-expressing cells were found in the malignant (3.5%), AT2 (2.2%) and club/secretory (2.4%) cell clusters (**Fig. 2a**). Among those clusters with > 1% *ACE2*-positive cells, AT2 cells expressed the highest expression of *ACE2* (**Fig. 2b**). We then explored expression patterns of other members of the RAS ^27^ (**Fig. 2c**). Of note, RAS genes were expressed at low levels in AT2 cells compared to other epithelial cell subsets and AT1 cells exhibited relatively higher expression of various RAS members such as adrenergic 2 beta receptor (*ADRB2*) (**Fig. 2c**). We then examined *ACE2* expression in subclusters of AT2 and malignant cells, the two populations with relatively highest fraction of cells positive for the SARS-CoV-2 receptor (**Fig. S2**). The AT2 subcluster (AT2_c2) with relatively highest expression of *ACE2* (**Fig. S2a** and **Fig. S2b**) exhibited similar levels of RAS genes compared to other AT2 subclusters except for elevated expression of amyloid precursor protein (*APP*) and the carboxypeptidase *CPM* (**Fig. S2c**). Similarly, we did not find similar patterns of expression between *ACE2* and other members of the RAS among the malignant cell subclusters (**Figs. S2c-e**). Earlier studies have shown that antihypertensive drugs such as losartan may impact expression levels of *ACE2* ^28, 29^. We thus analyzed *ACE2* expression in AT2 and lung malignant cells based on whether patients were on antihypertensive drugs. Interestingly, while the fractions of *ACE2*-positive AT2 and malignant cells were lower in losartan-treated patients, levels of *ACE2* expression in those cells were significantly higher in losartan treated patients (P < 10^−16^; **Fig. S3**). We also examined the expression and abundance patterns of *TMPRS22*, a serine protease recently shown to be crucial for SARS-CoV-2 spike protein priming upon host cell entry ^6^, as well as ADAM17, a sheddase that was shown to cleave (shed) ACE2 ^30^. *TMPRSS2* expression level and frequency were highest in AT2 cells among all lung subsets, whereas *ADAM17* expression levels were highest in malignant cells (**Fig. S4**). Also, among AT2 cells, expression levels of *TMPRSS2* and *ADAM17* were highest in the same AT2 subcluster as *ACE2* (**Figs. S2b and S4b**).

**Figure 2:**
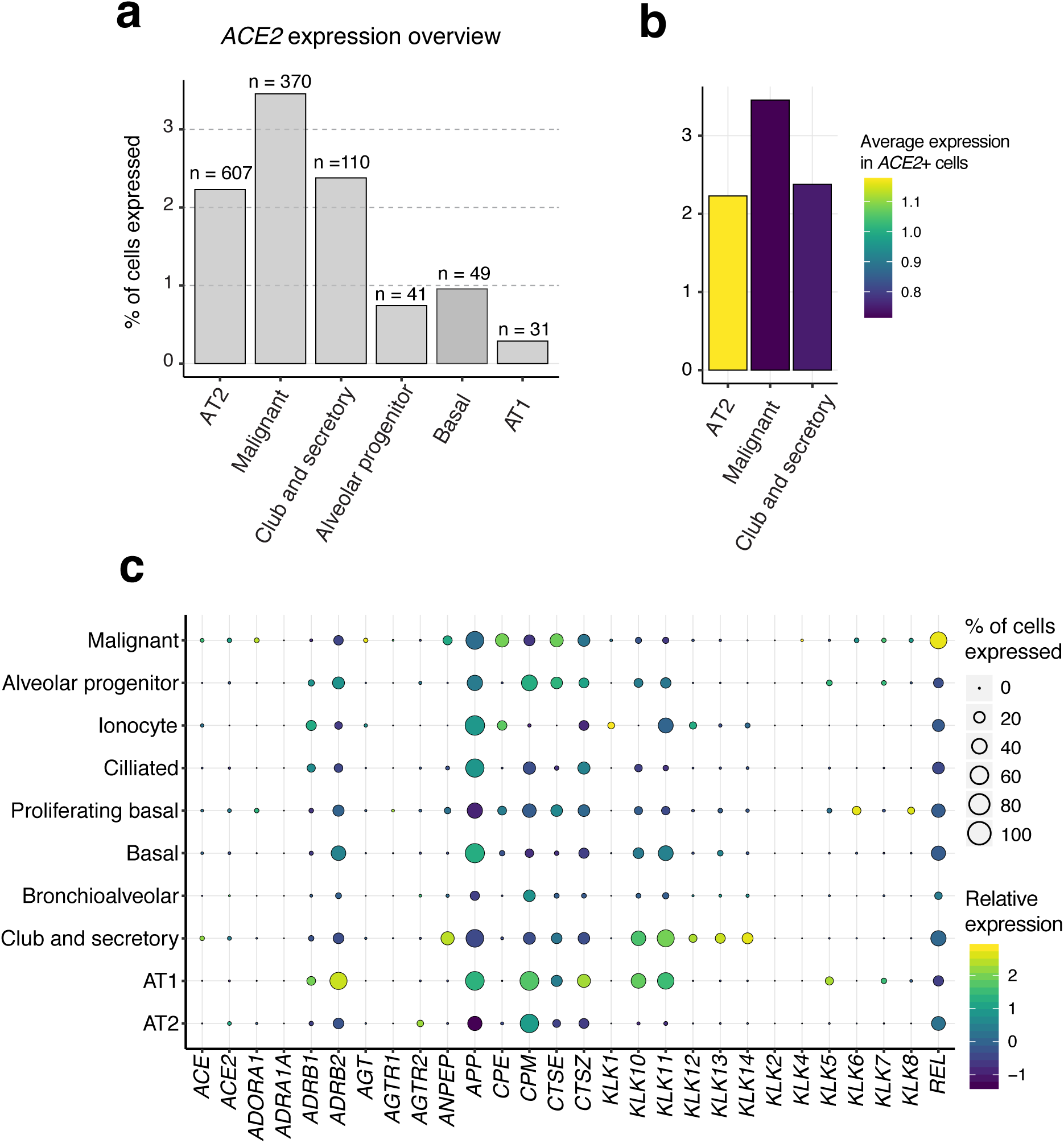
Single-cell expression analysis of *ACE2* and genes of the RAS in lung epithelium. **a)** Fraction of cells expressing the SARS-CoV-2 receptor *ACE2* in major airway lineage clusters. **b**) *ACE2* expression level (in *ACE2*-positive cells) among three airway lineage clusters with highest fractions of *ACE2*-expressing cells. **c**) Bubble plots showing cell fractions with scaled expression of genes of the RAS across the identified lung epithelial cell clusters.

We next sought to identify differentially expressed genes (DEGs) in AT2 cells based on *ACE2* expression. A cutoff of absolute gene expression fold-change >1.2 and an FDR q-value < 0.05 were applied to select DEGs between *ACE2*-expressing and *ACE2*-absent AT2 cells. We found genes up-regulated in *ACE2*-expressing AT2 cells that are highly pertinent to lung epithelial biology and disease such as *HHIP* (lung branching and chronic obstructive pulmonary disease/COPD, ^31, 32^), *FGG* (fibrosis, pneumonia and inflammation, ^33, 34^) and *C4BPA* (complement system, pneumonia and infection, ^35, 36^) (**Fig. 3**). In addition, we found that *ACE2*-expressing AT2 cells exhibited significantly higher expression of scavengers such as *CD36* ^37^ and *DMBT1*, a pattern recognition receptor that plays crucial host defense roles against bacterial and viral pathogens including influenza A and human immune deficiency virus I (HIV-I) ^38^. Interestingly, *DMBT1* expression was markedly and distinctly highest in AT2 cells relative to other lung cell subsets (**Fig. S5a**). Also, *DMBT1* expression in AT2 cells was higher compared to other coronavirus receptors such as the MERS-CoV receptor *DPP4* ^39, 40^ and *BSG* (also known as CD147) which was recently suggested to represent a potential alternate route of entry for SARS-CoV-2 ^41^ (**Fig. S5a**). Also, *DMBT1* displayed similar patterns of expression with *ACE2* among AT2 (**Fig. S5b**) and lung malignant cell (**Fig. S5c**) subclusters.

**Figure 3:**
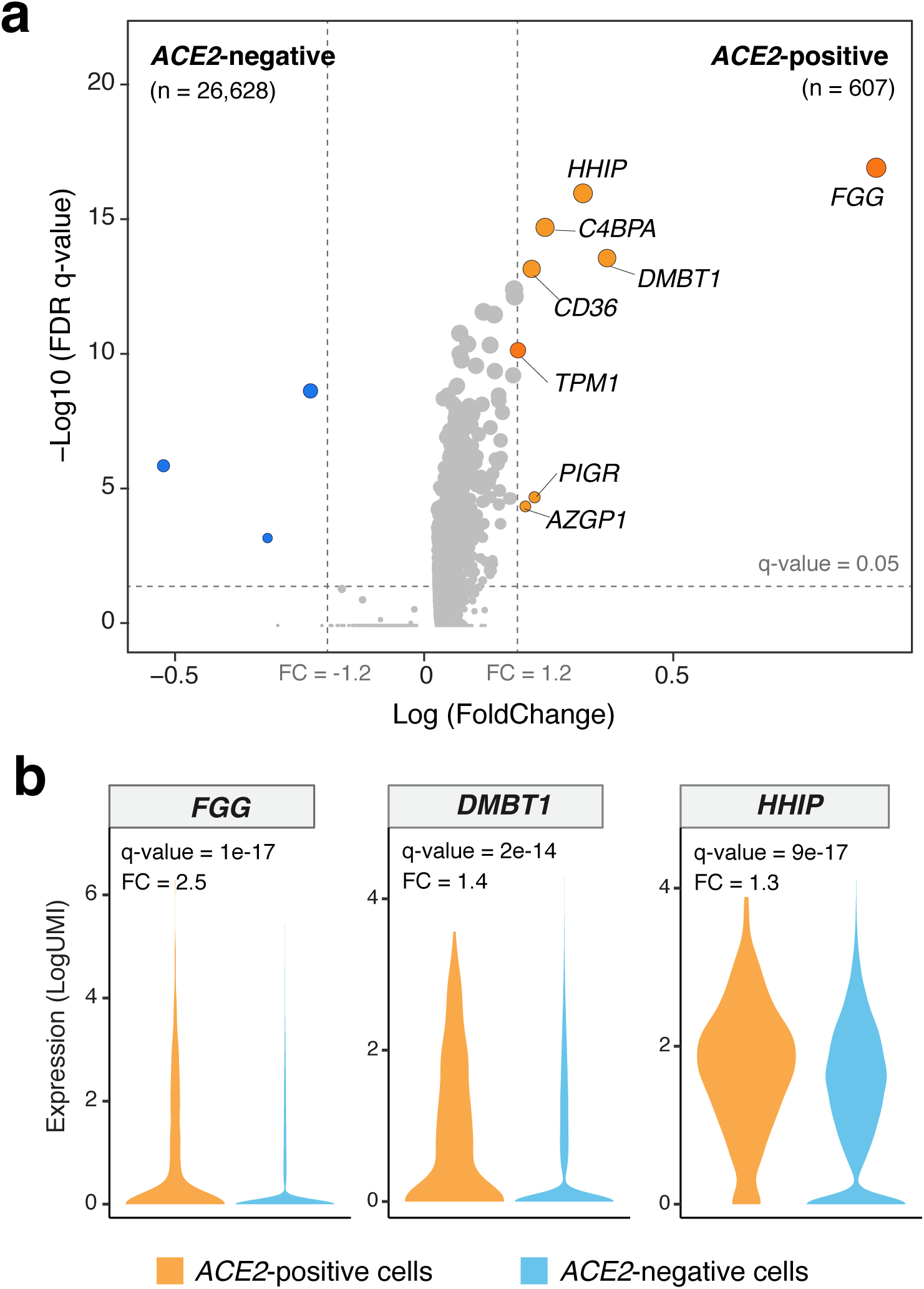
Differentially expressed genes between *ACE*2-expressing and -absent AT2 cells. **a**) Volcano plot showing significantly DEGs between ACE2-expressing (n = 607) an-absent (n = 26,628) AT2 cells. A cutoff of absolute gene expression fold-change >1.2 and an FDR q-value < 0.05 were applied to identify the DEGs. **b**) Violin plots showing significant up-regulation of *C4BPA, DMBT1* and *FGG* genes in *ACE2*-expressing compared with -absent AT2 cells, FC: Fold change.

Recently, expansion of ACE2-expressing lung epithelia in response to cigarette smoke has also been proposed ^42^ and smoking was shown to activate the RAS ^43^. We were thus enticed to compare the identified *ACE2*-co-expressed genes, along with members of the RAS, in lung epithelial cells based on patient smoking status. We identified genes differentially expressed between AT2 cells derived from smokers (at least 20 pack year smoking history, n = 10,901 cells) and those from non- or light-smokers (less than five pack years, n = 16,334 cells) based on a q-value threshold of 0.05 and a fold-change cut-off of 1.5 (**Fig. 4a**). While *ACE2* itself was not modulated by smoking status, we found that the virus receptor and scavenger *DMBT1*, as well as *FGG* and *C4BPA*, were significantly increased in smoker compared to non/light smoker AT2 cells (**Fig. 4a-b**). *DMBT1* was not significantly differentially modulated among malignant cells based on smoking status (**Fig. 4c**), though we found increased expression of *BSG* and members of the RAS (mostly kallikreins) in smoker (n = 9,047) compared to non/light smoker (n = 1,657) malignant cells (**Fig. 4c-d**).

**Figure 4:**
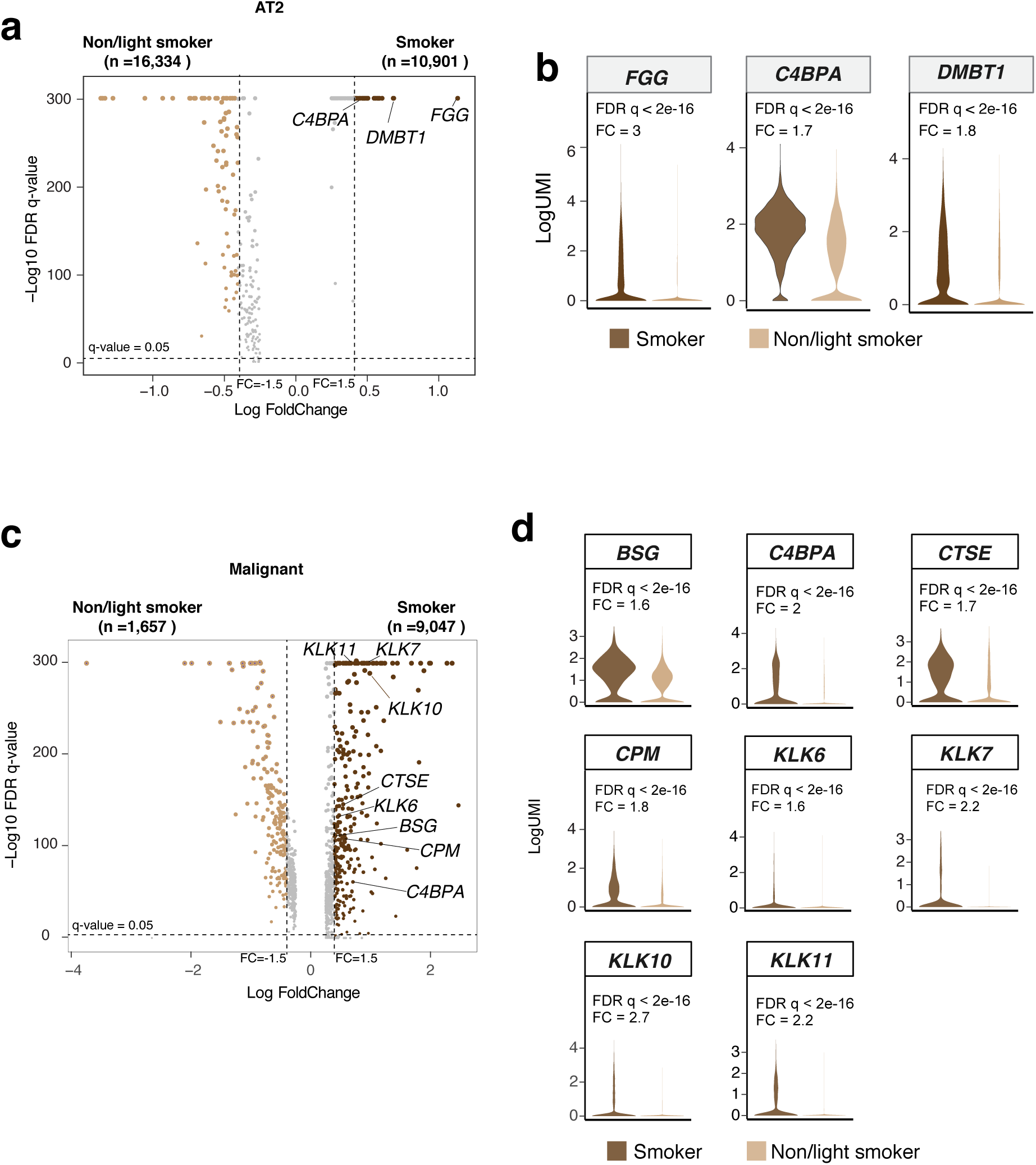
Differentially expressed genes between AT2 and lung malignant cells from smoker and non/light smoker patients. **a**) Volcano plot showing significantly DEGs between AT2 cells from smoker (n = 10,901 cells) and non/light smoker (n = 16,334 cells) patients. **b**) Violin plot showing significant upregulation of *FGG, C4BPA* and *DMBT1* in smoker AT2 cells. **c**) Volcano plot showing significantly DEGs between malignant cells from smoker (n = 9,047 cells) and non/light smoker (n = 1,657) patients. **d**) Violin plots of select significantly up-regulated genes in smoker malignant cells. A cutoff of absolute gene expression fold-change >1.2 and an FDR q-value < 0.05 were applied to identify the DEGs. FDR q-values < 1× 10^−300^ were trimmed to 1 × 10^−300^ for graphical purposes.

Our findings on the significant associations between *ACE2* and *DMBT1* prompted us to probe the correlation of both genes in the AT2 compartment. We performed pseudo-bulk analysis of the AT2 cluster (by sample) in our cohort and found that *DMBT1* and *ACE2* exhibited positive correlation (R = 0.3) albeit not reaching statistical significance (**Fig. 5a**). To validate this finding in larger cohorts, we performed deconvolution analysis of bulk RNA-seq data of the cancer genome atlas (TCGA) LUAD cohort to estimate the abundance of AT2 cells in normal lung tissues (n = 110). Both *DMBT1* and *ACE2* expression levels significantly and positively correlated with AT2 fractions (indicated by the AT2 meta-score, *P* < 0.05) (**Fig. 5b**). Interestingly, *BSG*, the other recently proposed entry point for SARS-CoV-2, also significantly and positively correlated with the AT2 meta-score in the normal lung tissues (*P* < 0.05) (**Fig. 5b**). Similar results were obtained with *TMPRSS2* (*P* < 0.05) but not *ADAM17* (**Fig. S6a**). Further, among TCGA normal lung tissues with high AT2 fractions (meta-score > 15.47), *DMBT1* expression significantly and positively correlated with *ACE2* (R = 0.41, *P* < 0.05) (**Fig. 5c**). We did not find significant correlation between *ACE2* and *TMPRSS2* nor with *ADAM17* in the TCGA lung tissues by deconvolution analysis (**Fig. S6b**). Our findings point to specific *ACE2*-expressing AT2 cells that also co-express other pathogen receptors and scavengers (e.g. *DMBT1*) thus possibly representing a minute subpopulation in the lung with unique host defense properties and functions.

**Figure 5:**
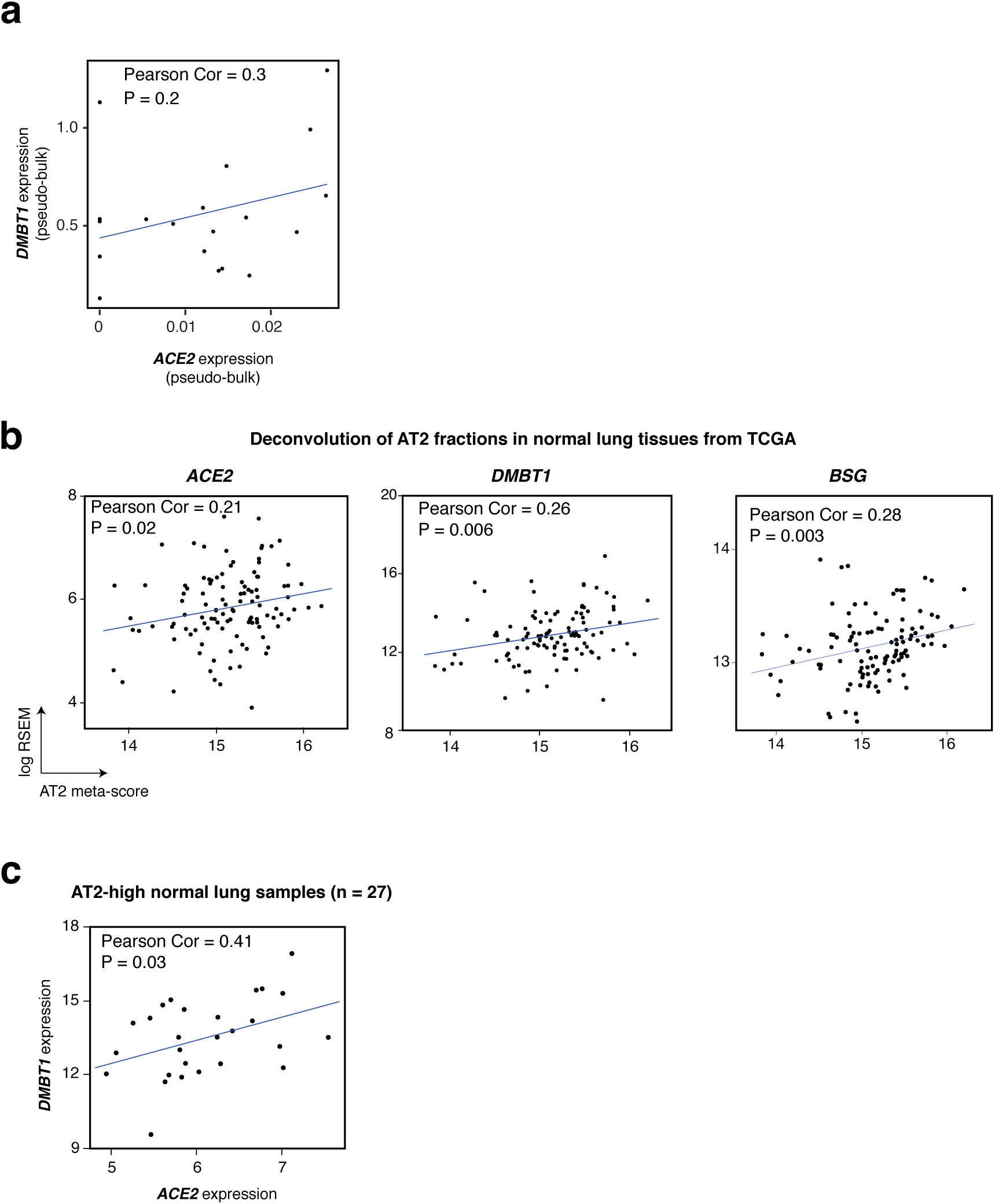
Expression of the viral scavenger *DMBT1* correlates with that of *ACE2* in lung AT2 cells. **a**) Correlation between *ACE2* and *DMBT1* expression in pseudo-bulk data from this study. **b**) Scatter plots showing correlations between estimated AT2 cell fractions and *ACE2, DMBT1* and *BSG* in TCGA normal lung samples. **c**) Significant correlation between *ACE2* and *DMBT1* in TCGA normal lung samples with high AT2 cell fractions (meta-score > 15.47). Correlations were statistically analyzed using Pearson’s correlation coefficient.

## DISCUSSION

We interrogated our relatively large in-house single-cell RNA-sequencing dataset of 70,085 lung epithelial cells to examine abundance and expression patterns of the SARS-CoV-2 receptor *ACE2*. While *ACE2* was expressed in a low fraction of lung epithelial cells (roughly 1.7%), its levels among all lung cell subsets were highest in AT2 cells. *ACE2*-expressing AT2 cells co-expressed genes with crucial roles in lung pathological conditions such as COPD, pneumonia and pathogen infection including the viral binding scavenger *DMBT1* and which overall were higher in cells derived from smokers relative to non/light smokers. We also found that *DMBT1* positively correlated with AT2 cell fractions and with *ACE2* itself within the AT2 compartment. Our findings suggest that *ACE2*-expressing cells in the lung are relatively scarce, they exhibit unique molecular and biological features that are pertinent to antiviral and host defense functions by the lung epithelium.

Our findings on expression of *ACE2* in AT2 cells are in line with previous studies ^44, 45^. Hamming et al showed abundant immunohistochemical expression of ACE2 protein in AT2 cells in the lung ^45^. *Ace2* was shown to not only to be expressed in AT2 cells in the mouse lung but also to regulate alveolar epithelial cell survival ^46^. Of note, Mossel and colleagues demonstrated that SARS-CoV, by binding to ACE2, replicates primarily in AT2 and not AT1 cells ^44^. Also, AT2 cells were shown to be primarily responsible in the lung for an innate immune response against infection with SARS-CoV ^47^. Interestingly, a very recent preprint by Zhao and colleagues revealed that *ACE2* expression was concentrated in a small population (∼1.4%) of AT2 cells ^48^. Our findings are in very close agreement with the preprint by Zhao et al in which we find that the SARS-CoV-2 receptor *ACE2* is expressed in a minute fraction of AT2 cells (∼2.2%). Here, we further interrogated a relatively larger and more diverse repertoire of airway lineages, from both normal and lung tumor (LUAD) tissues, and found that *ACE2* expression was not only highest in AT2 cells but also that malignant cells exhibited the highest *ACE2*-positive fraction. It is worthwhile to mention that patients with cancer, including lung malignancy, may represent a population that is extremely vulnerable to COVID-19 ^49^. Our finding on expression of *ACE2* in lung malignant (adenocarcinoma) cells may have implications for the management of COVID-19 in lung cancer and clinical studies are warranted to explore this conjecture. Our analysis also revealed significantly elevated *ACE2* levels, among *ACE2*-positive AT2 and malignant cells, in patients who were on hypertension treatment with losartan compared to those without antihypertensive therapy. While our finding is in line with previous reports suggesting that antihypertensive agents may augment *ACE2* expression ^28, 29, 50^, it needs to be interpreted with caution. Our patient cohort is very limited in size and our study was designed with a focus on single-cell analysis of lung epithelial cells. We thus cannot construe the implications of this finding on COVID-19 management ^50, 51^ and further studies are warranted to better (e.g., larger cohort) explore the association between treatment with antihypertensive drugs and *ACE2* expression in human cells.

We found that *ACE2*-positive compared to *ACE2*-negative AT2 cells exhibited increased levels of genes with crucial roles and expression features in lung pathological diseases. The hedgehog interacting protein *HHIP* was not only shown to play important roles in airway branching during lung development ^32^, but also single nucleotide polymorphisms of this gene are associated with increased risk for COPD ^31^, a pulmonary ailment characterized by chronic inflammation ^52^. *FGG* coding for fibrinogen-gamma was shown to be induced by pro-inflammatory cytokines ^34^ and to be elevated in lung pneumonia and infection ^33^. *C4BPA*, coding for C4BP and part of the complement system, was found to recognize and bind pneumonia-causing streptococci in the lung epithelium ^35, 36^. It is noteworthy that many patients with COVID-19 (e.g. those with severe disease) commonly display the same pathological phenotypes, namely lung inflammation, fibrosis, and pneumonia, linked to those *ACE2-*co-expressed genes. It is intriguing to suggest that perhaps this small population of *ACE2*-expressing cells may underlie the pathogenesis of severe acute respiratory distress syndrome, pneumonia and respiratory failure in COVID-19 patients. It is important to note that emerging studies are showing that males are disproportionately affected by COVID-19, possibly due to sex-based immunological bias or differences in smoking patterns and prevalence ^53^. Also, a recent preprint suggested that smoking causes expansion of *ACE2*-expressing lung epithelia ^42^. Interestingly, several of the *ACE2*-co-expressed genes we identified (*FGG, DMBT1, C4BPA*) were elevated in smoker AT2 cells. Of note, smoking increases lung inflammation and the risk for various lung diseases including interstitial lung fibrosis, pneumonia and COPD ^54^. It is plausible that smoking may augment SARS-CoV-2 entry in the lung and COVID-19 pathogenesis, a supposition that warrants future studies.

A notable finding in our study was the co-expression of pathogen including viral scavengers and receptors such as *CD36* and *DMBT1* in *ACE2*-positive AT2 cells. We also found that among all lung subsets, AT2 cells distinctly and markedly displayed the highest expression of *DMBT1*. Further, *DMBT1* correlated with AT2 fractions in independent cohorts of bulk-sequenced lung tissues and positive correlated with *ACE2* in the AT2 compartment. DMBT1, also known as gp340, was shown to inhibit influenza A by binding to hemagglutinin (HA) on the virus ^55^. DMBT1 was also shown to interact with surfactant protein D (SFTPD) in alveolar cells to agglutinate and inhibit influenza A virus ^38^. It has been suggested that the antiviral drug oseltamivir cooperates with innate immune proteins such as DMBT1 in inhibition of lung epithelial cell infection by influenza A virus ^55^. Interestingly soluble DMBT1 in saliva was shown to exert host defense roles (neutralization or inhibition of oral transmission) against influenza A virus ^55^ as well as HIV-1 ^56^. DMBT1 was shown to specifically inhibit HIV-1 infectivity by binding to the virus envelope protein gp120, the same protein that binds to the CD4 receptor on T cells in the host ^57^. Intriguingly, very much like emerging reports on ACE2 acting as an entry route in the lung that facilitates SARS-CoV-2 infection ^6^, DMBT1 was shown to aid in HIV-1 transcytosis across genital tract tissue ^58^. Also, the study by Stoddard and colleagues demonstrated that transport of HIV-1 can be inhibited by antibodies or peptides that block the interaction of DMBT1 with the HIV-1 envelope protein gp120 ^58^. Given our findings on *DMBT1* expression in lung AT2 cells and its co-expression with *ACE2*, as well as its reported binding with multiple viruses, we suggest that targeting DMBT1, whether by adding soluble DMBT1-derive peptides ^59^ or by antibody-based neutralization, may represent a viable strategy to counteract SARS-CoV-2 infection and ameliorate COVID-19.

In conclusion, single-cell transcriptomic analysis of our lung epithelial cell cohort demonstrated that among all lung cell subsets AT2 cells displayed highest relative expression of the SARS-CoV-2 receptor *ACE2*, albeit at a low cell fraction (∼2.2% of all AT2 cells). We found that the viral scavenger *DMBT1* is highly expressed in AT2 cells and correlates with *ACE2* in this compartment. Our study points to a scarce *ACE2*-expressing AT2 population that also expresses genes involved in inflammatory lung pathological conditions (e.g., pneumonia) and in host defense and, thus, could be exploited (or repurposed) as phenotypic targets for treatment of SARS-CoV-2 infection and COVID-19 disease.

---

**Figure S1.**
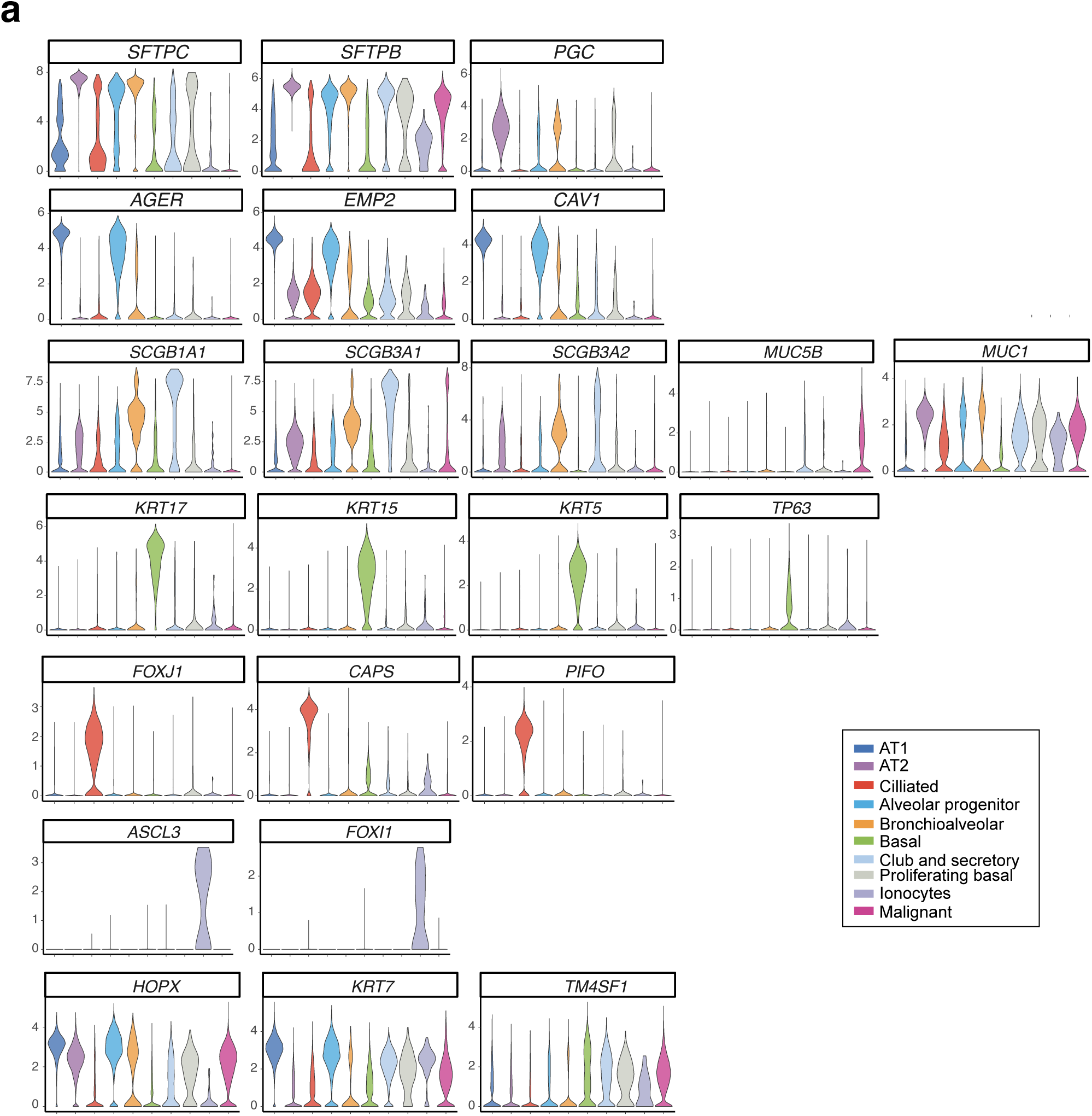

**Figure S2.**
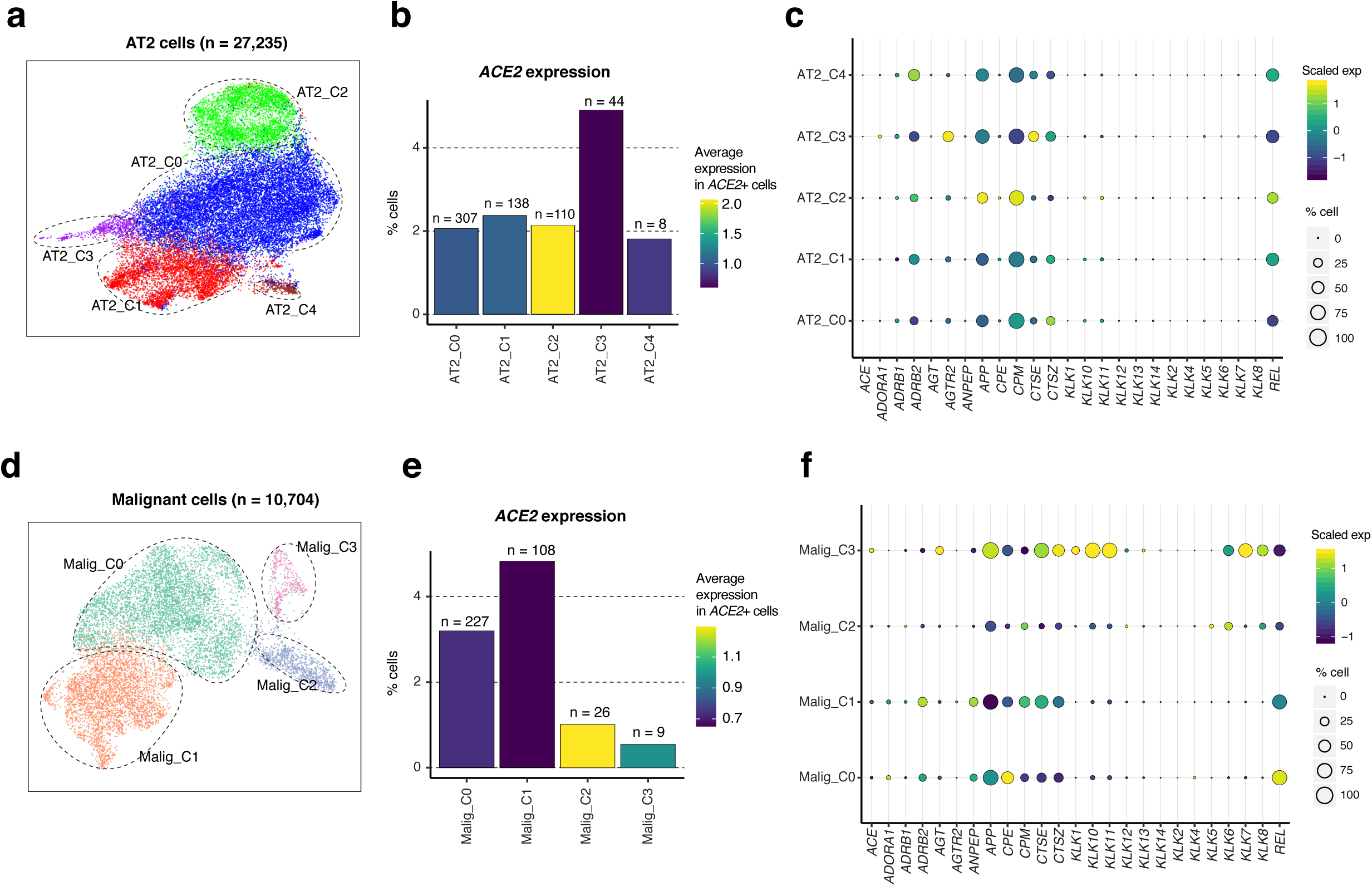

**Figure S3.**
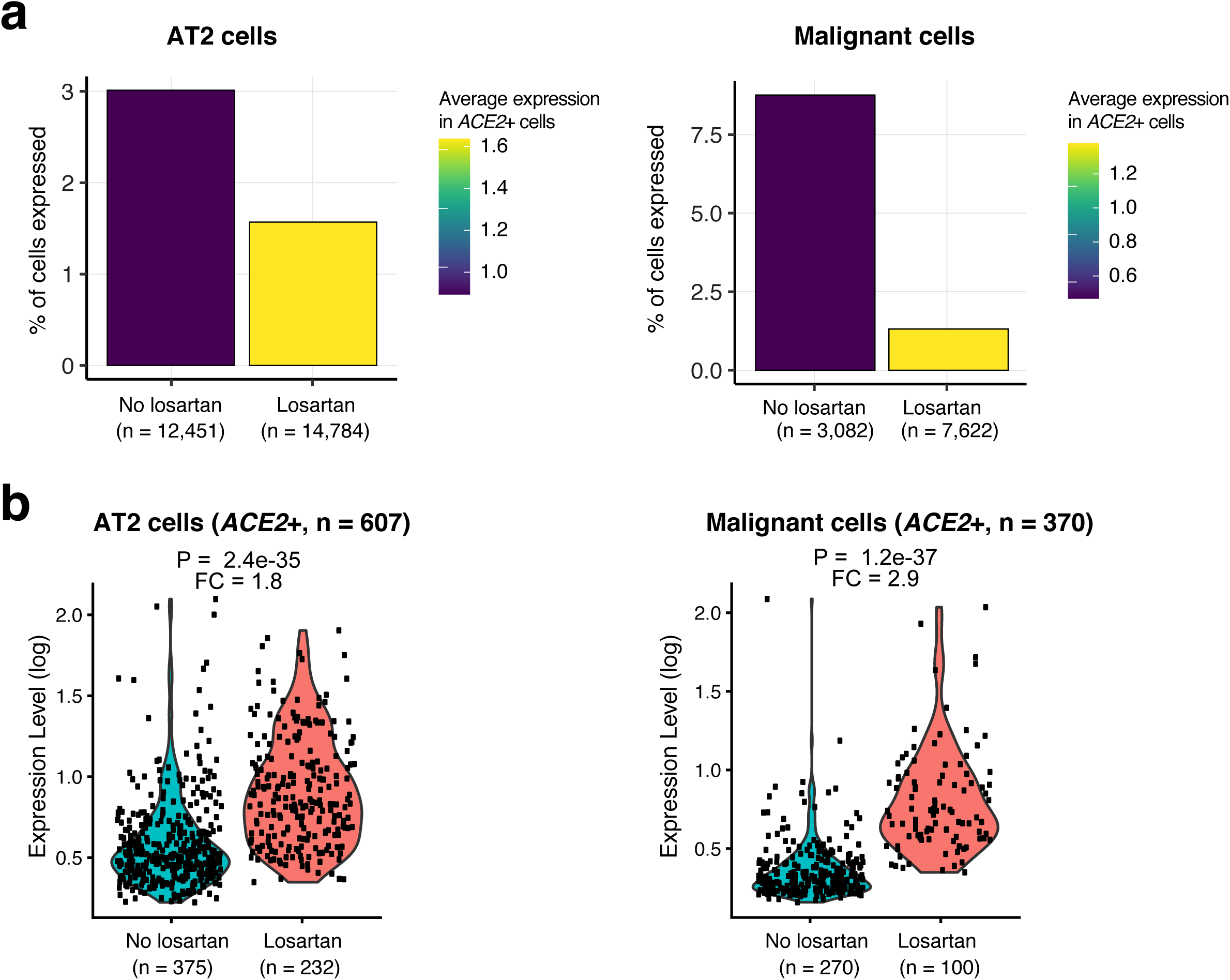

**Figure S4.**
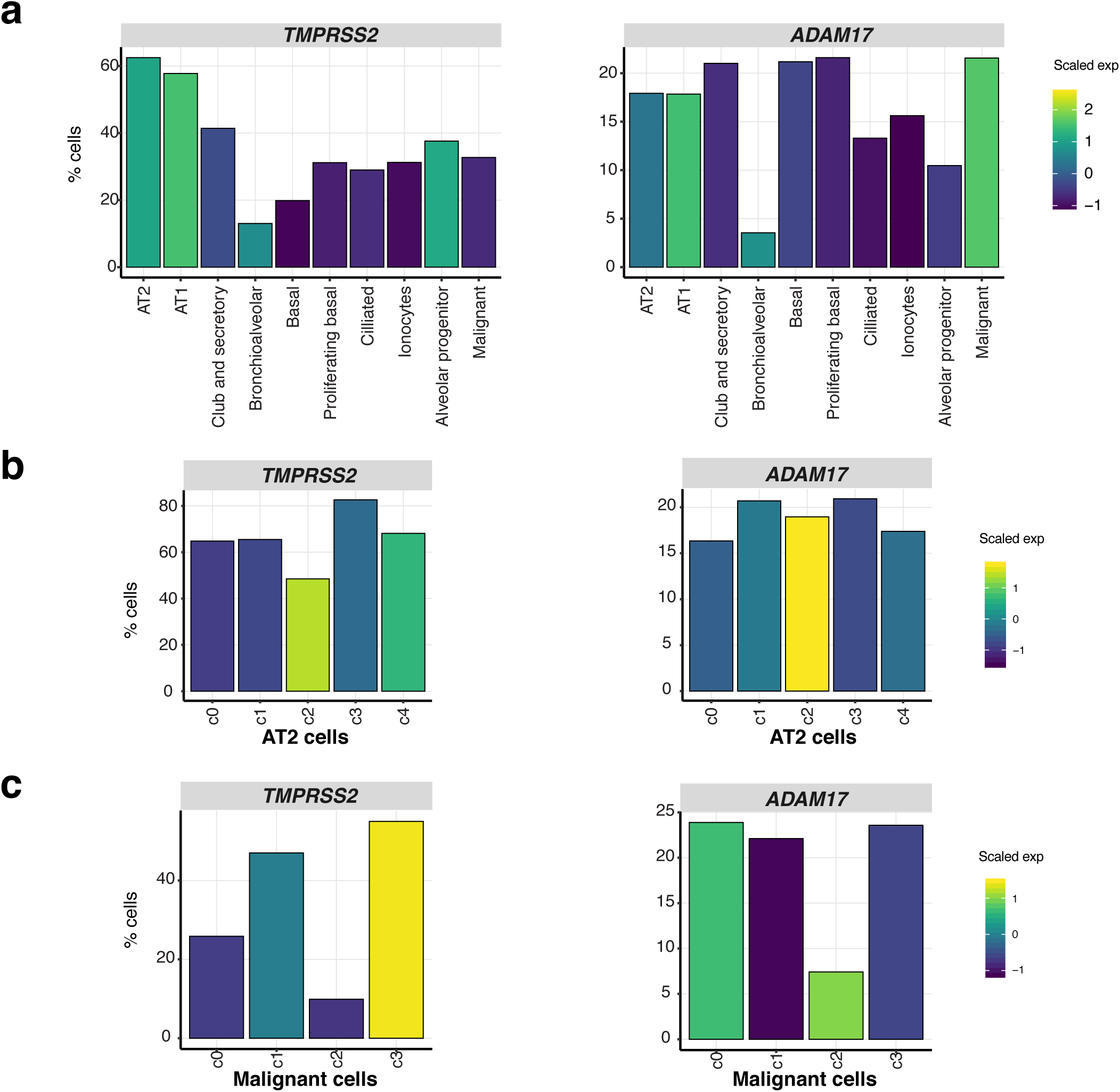

**Figure S5.**
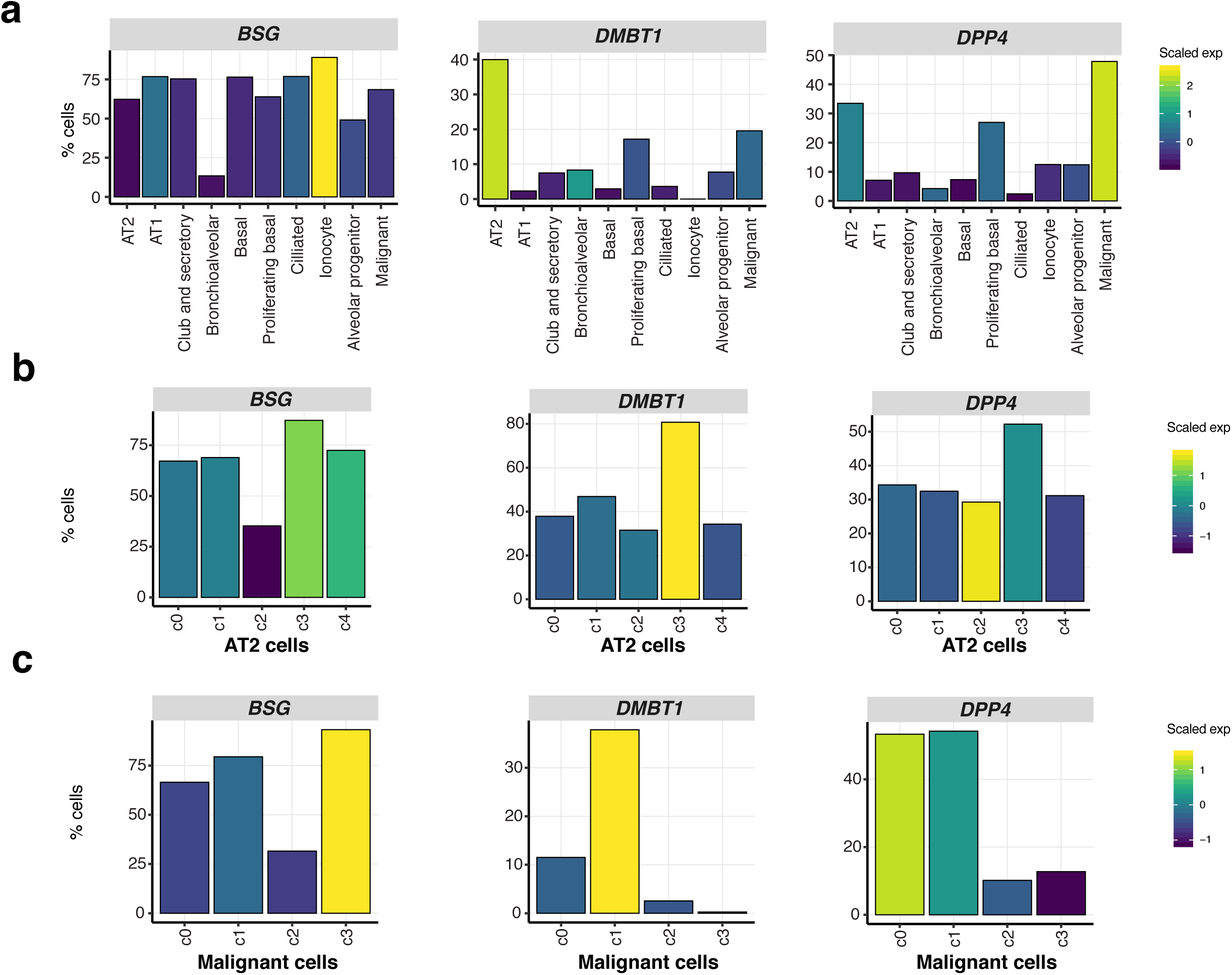

**Figure S6.**
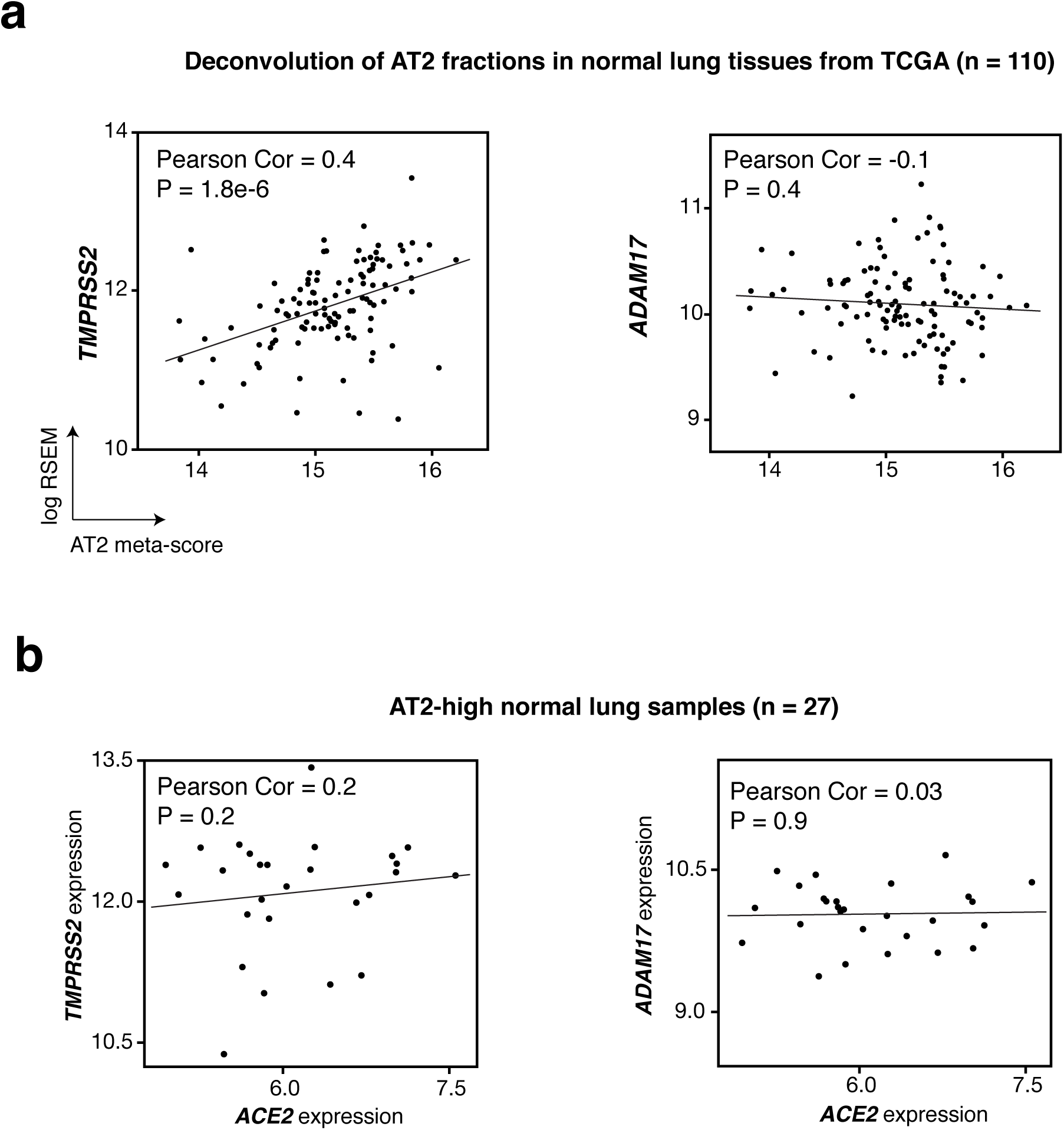

