## Supplementary figure legends for "Single-cell analysis of human lung epithelia reveals concomitant expression of the SARS-CoV-2 receptor ACE2 with multiple virus receptors and scavengers in alveolar type II cells"

**Figure S1: Expression of airway lineage markers in the identified lung epithelial clusters.** Violin plots showing differential expression of major airway cell lineage markers among the identified 10 lung epithelial cell clusters.

**Figure S2: Expression analysis of *ACE2* and genes of the RAS in AT2 cells and lung malignant cell sub-clusters.** UMAP plots showing AT2 (**a**) and malignant (**d**) cell sub-clusters. *ACE2* expression frequency and level among AT2 (**b**) and malignant (**e**) cell subclusters. Bubble plots showing cell fractions with scaled expression of genes of the RAS among AT2 (**c**) and malignant (**f**) cell sub-clusters.

**Figure S3: Expression analysis of *ACE2* in AT2 cells and lung malignant cells based on treatment with losartan. a**) Expression frequency and levels of *ACE2* in *ACE2*-positive AT2 (left) and lung malignant cells (right) between patients with and without losartan treatment. **b**) Violin plots showing significant up-regulation of *ACE2* genes in *ACE2*-expressing AT2 and malignant cells from patients who received losartan relative to those who did not receive antihypertensive treatment, FC: Fold change.

**Figure S4: Expression analysis of the proteases *TMPRSS2* and *ADAM17* in lung epithelial cells. a)** Expression frequency and levels of *TMPRSS2 and ADAM17* in the 10 identified lung epithelial cell clusters. Expression frequency and levels of *TMPRSS2* and *ADAM17* among AT2 (**b**) and malignant (**c**) cell subclusters.

**Figure S5: Expression analysis of *DMBT1* and the coronavirus receptors *BSG* and *DPP4* in lung epithelial cells. a)** Expression frequency and levels of *BSG*, *DMBT1* and *DPP4* in the 10 identified lung epithelial cell clusters. Expression frequency and levels of *BSG*, *DMBT1* and *DPP4* among AT2 (**b**) and malignant (**c**) cell subclusters.

**Figure S6: Analysis of correlation of *TMPRSS2* and *ADAM17* expression with *ACE2* in lung AT2 cells. a**) Scatter plots showing correlations of estimated AT2 cell fractions with *TMPRSS2* and *ADAM17* in TCGA normal lung samples. **b**) Scatter plots showing correlations of *ACE2* expression with *TMPRSS2* and *ADAM17* levels in TCGA normal lung samples with high AT2 cell fractions (meta-score > 15.47). Correlations were statistically analyzed using Pearson's correlation coefficient.
